# A genetic screen to identify host factors required for commensal bacterial adherence to the gut epithelia

**DOI:** 10.64898/2026.11.09.765429

**Authors:** Anupama Singh, Dalaena E Rivera, Cayla Cruse, Truc Nguyen, Priya Gupta, Kayla Poirier, Robert J Luallen

**Affiliations:** Department of Biology, San Diego State University, San Diego, CA 92182, USA

## Abstract

Bacterial adherence to gut epithelial cells likely involves host genetic components. We isolated a gut commensal bacterium, *Lelliottia jeotgali*, that can robustly colonize *C. elegans* through attachment to intestinal epithelial cells. Here, we developed a novel screen in *C. elegans* to identify host genes required for microbiome bacterial colonization. Using transgenic *L. jeotgali* strain expressing tdTomato, we conducted forward genetic screens in *C. elegans* to select for mutants that completely lost colonization. We identified multiple allelic mutations in five different *C. elegans* genes, termed <u>k</u>nockout <u>o</u>f *<u>L</u>elliottia* <u>a</u>dherence *(kola)* mutants, with a near complete loss of adherence. Each of *kola-1-5* is uncharacterized and predicted to encode for a secreted or membrane-bound protein. We also identified *gly-5*, a GalNAc transferase that initiates O-linked glycosylation. Overall, we describe the first *C. elegans* screen to identify host factors that control microbiome bacterial colonization in the gut.

## Introduction

The capacity for microbes to colonize a host are controlled by many host genes that play a role in facilitating or blocking microbial colonization, retention, and niche formation (Wilde et al. 2024). Genetic screens in diverse model systems have identified microbial determinants of host colonization (Kline et al. 2009; Lee et al. 2013; Powell et al. 2016). Forward genetic screens in *C. elegans* that have identified host factors affecting pathogen colonization, often hitting host immune regulators (Hodgkin et al. 2000; Marsh and May 2012; Tsai et al. 2021). Similar screens using gut microbiome bacteria are limited, likely due to the sheer quantity of bacteria in many microbiomes, the complexity of host mucosal surfaces, and difficulty in visualizing colonization in many animals (Linden et al. 2008; Vonaesch et al. 2018). Additionally, many microbiome bacteria interact within complex mucosal niches that require navigating processes like attachment and feeding off mucus layers, avoiding antimicrobial factors, and outcompeting other microbes, making screens potentially confounding (Dishaw et al. 2024). Here, we describe the first genome-wide screen to identify host factors required for commensal microbiome colonization in *C. elegans*, using a wild-isolated commensal bacteria, *Lelliottia jeotgali*, that attaches to intestinal epithelial cells in the lumen (Rivera et al. 2025).

*C. elegans* has emerged as a promising model system to study host-microbiome interactions (Singh and Luallen 2024). The recent extensive worldwide sampling of wild *Caenorhabditis* nematode populations from rotten fruits and plant matter, has led to the isolation and identification of its naturally associated microbiota, including numerous bacterial species colonizing the *C. elegans* gut (Félix and Duveau 2012; Dirksen et al. 2016). But not many *C. elegans* commensals are known to colonize the gut robustly, unless the population of animals are forced to continuously consume the commensal bacteria (Rivera et al. 2025). We discovered that *L. jeotgali* can robustly colonize lab-grown, wild-type *C. elegans* through attachment to intestinal epithelial cells in the lumen with no net negative fitness effect on the host (Rivera et al. 2025). Additionally, this commensal bacterium can actively replicate in the gut lumen, as a brief exposure of *C. elegans* to *L. jeotgali* results in nearly 100% of animals being fully colonized across the anterior-posterior (A-P) length of the gut. These facts make *L. jeotgali* an ideal commensal microbiome bacterium to search for host genes mediating microbiome bacterial colonization.

Here, we present an unbiased forward genetic EMS screen in *C. elegans* to identify factors that are required for *L. jeotgali* colonization of the gut. For easy visualization of colonization, we created transgenic *L. jeotgali* strain expressing tdTomato using transposon insertion into the bacterial genome.

The screen identified over 40 mutants with a near complete loss of *L. jeotgali* colonization. Using in silico complementation, we found these mutants strains had alleles in five different uncharacterized genes, which we named <u>k</u>nockout <u>o</u>f *<u>L</u>elliottia* <u>a</u>dherence *(kola)*. We also identified *gly-5*, a GalNAc transferase predicted to initiate O-linked glycosylation in the intestine (Hagen and Nehrke 1998). Overall, we describe the first *C. elegans* screen to identify host factors that control microbiome bacterial colonization in the gut.

## Materials and Methods

### Strains

Wild type *C. elegans* N2 strain was used for mutagenesis. VC3230 *gly-5*(*gk3119*) III was acquired from the *C. elegans* Genetics Center. ERT413 jySi21[*spp-5p::GFP; cb-unc-119(+*)] II was obtained from the Troemel lab (Luallen et al. 2016). Worms were maintained on standard Nematode Growth Media (NGM) agar plates seeded with strain OP50-1 *E. coli*. For the microbiome bacteria strain LUAb3 of *L. jeotgali* was isolated from a wild *C. elegans* nematode sampled from the SDSU campus (Rivera et al. 2025). Transgenic *L. jeotgali* strain expressing tdTomato (LUAb18) was generated using transposon insertion into the bacterial genome. LUAb3 and LUAb18 were grown in standard LB broth and LB plates in vitro at 32°C.

### Transposon insertion

To insert a tdTomato fluorescent tag into LUAb3, we utilized mini-Tn7 transposons, as previously described (Lambertsen et al. 2004; Tran et al. 2022). Briefly, we transformed a streptomycin resistant LUAb3 strain with mini-Tn7 delivery plasmids harboring a tdTomato tag via conjugation. Successful insertion of the tag was validated by whole genome sequencing.

### EMS mutagenesis screen for loss of microbiome colonization

To conduct the forward genetic screen, *C. elegans* N2 were bleached and grown on Nematode Growth Media (NGM) plates seeded with *E. coli* OP50 until they reached the L4 stage. Animals were harvested and washed once in M9. The population was split into 2 x 15 mL conical tubes and resuspended in 4 mL of M9. Into one tube, 15 µl of EMS (bought as 10M stock) was added to the 4 mL of M9 to reach a concentration of 37.5 mM. The second tube received no EMS and served as a negative control. Both tubes were capped, parafilmed, and incubated at room temperature on a rotator for 4 h. To remove the EMS, the tubes were spun down and the supernatant was removed using a Pasteur pipet. Both the supernatant and pipet were disposed of into a 50 mL tube containing NaOH. The tubes were washed twice with 5 mL of M9, using the same disposal method into NaOH. About 30 animals in 500 µl of 10x concentrated *E. coli* OP50 were placed on a 10 cm NGM plate seeded with OP50-1. Each plate is considered a “pool”. About 3-7 pools were generated for each screen, with a total of 3 independent screens conducted. The parental (P0) generation was incubated at 20°C for 5 days to give rise to the F1 generation. Once the F1 generation became gravid adults, they were bleached and synchronized. The resulting F2 L1s were plated onto NGM seeded with LUAb3-tdTomato. After 2 days, the F2 generation was screened for loss of colonization on a fluorescence-capable Nikon SMX-18 stereoscope. To help with the screening process, these animals were harvested, washed 2-3 times in M9 to get rid of external bacteria, and plated onto OP50 seeded 10 cm NGM plates. As the animals defecate, transient bacteria in the intestine was removed. Animals that have lost red fluorescence were picked onto their own 3.5 cm NGM plate seeded with OP50 for propagation and further validation of the phenotype. These mutants were given the phenotypic designation **A**dhering **b**acteria **c**olonization **d**eficient (Abcd).

### Whole genome sequencing and in silico complementation

To identify genes responsible for Abcd phenotypes, we conducted whole genome sequencing and in silico complementation to identify genes with multiple allelic hits across the validated Abcd mutant strains. Our lab N2 strain and the Abcd mutant strains were sequenced by Novogene on an Illumina NovaSeq X Plus platform, 150 bp paired-end reads. Reads were aligned to the *C. elegans* reference genome (WBcel235; GCF_000002985.6) using BWA-MEM (v0.7.17-r1188; parameters: mem - t 4 -k 32 -M) (Li and Durbin 2009). Reads were sorted and duplicates merged with SAMtools (v1.13) and Picard (v1.111) (Li et al. 2009). Single nucleotide variants (SNVs) were called using SAMtools mpileup (v1.3.1; parameters: -m 2 -F 0.002 -q 1 -C 50 -t SP - t DP -t AD), and filtered to retain variants with more than 4 and fewer than 10^7^ reads and mapping quality (MQ) greater than 20. Structural variants (SVs) were identified using BreakDancer (v1.4.4; parameter: -q 20) based on discordant paired-end read mapping and insert-size distributions, retaining calls supported by at least 2 paired-end reads (Chen et al. 2009). Functional annotation of SNVs, InDels, and SVs (insertions, deletions, and inversions) was performed with ANNOVAR (2015Dec14 release) (Wang et al. 2010).

We used in silico complementation to identify identical genes mutated across multiple Abcd mutant strains as described (Minevich et al. 2012). Briefly, in a database containing all SNVs across every sequenced strain, all background variants were removed, defined as homozygous variants found in our N2 strain or found in >75% of Abcd strains. Then, we searched for genes containing either (i) at least three different moderate-(missense) or high-impact (splice site or nonsense) homozygous variants, each in a different Abcd mutant strain, or (ii) at least two different high-impact homozygous variants, each in a different Abcd mutant strain.

### Backcrossing, genotyping, and complementation

Hermaphrodites for multiple *kola* strains were backcrossed four times to N2 males, in cluding four alleles of *kola-1*, six alleles of *kola-2*, both alleles of *kola-3*, four alleles of *kola-4*, and three alleles of *kola-5*. Males were produced using heat shock of 4-6 L4 hermaphrodites at 32°C for 2-4 hrs. Progeny were phenotyped for a gain of colonization at the F_1_ generation and loss of colonization at the F_2_. Then, F_2_ animals were genotyped for the corresponding *kola* mutation using PCR amplification with primers flanking the mutation (Table 1) and Sanger sequencing of the amplicon. This process was repeated for a total of four backcrosses.

**Table 1.**
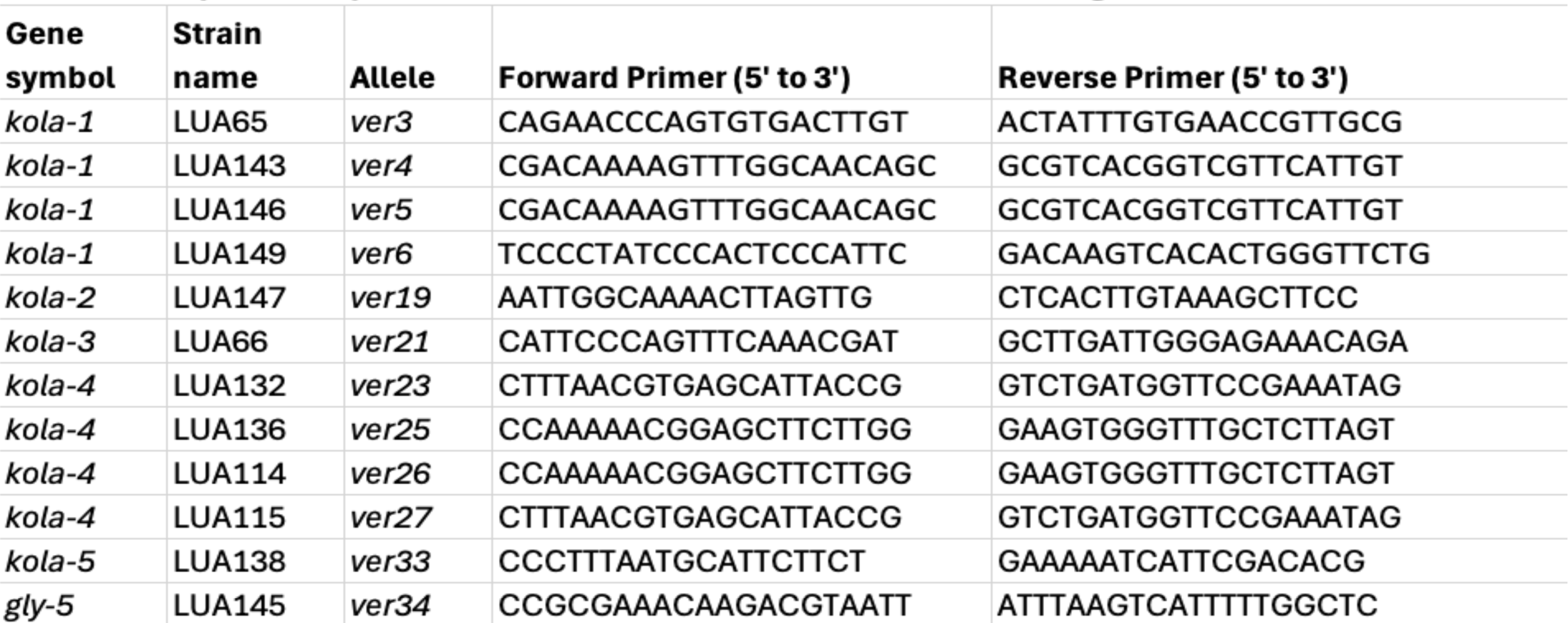
PCR primer sequences used to validate alleles after backcrossing.

Traditional complementation was used to further validate *kola-1, kola-3* and *kola-4*, with strains carrying two unique alleles of each gene failing to rescue an Abcd phenotype at the F_1_ generation. As a control, we performed complementation tests between different *kola* mutants, resulting in loss of Abcd phenotypes. For *gly-5 (ver34)*, we obtained another allele, *gly-5*(*gk3119*) and performed an indirect complementation test using outcrossed heterozygous males. For this, N2 males were crossed into *gly-5(ver34)* (P_0_) to produced heterozygous *gly-5(ver34)* males (F_1_). These males were crossed into *gly-5(gk3119)* to test for complementation in *ver34/gk3119* trans-heterozygous F_2_ males.

### Phenotyping for microbiome colonization assays

*L. jeotgali* colonization of N2 and Abcd strains was conducted as described (Rivera et al. 2025). Briefly, an overnight culture of LUAb3 (*L. jeotgali*) or LUAb18 (*L. jeotgali-*tdTomato) was grown in LB broth at 32°C prior colonization. Bleached and synchronized *C. elegans* were grown on standard NGM plates seeded with *E. coli* OP50-1 to the L4 stage. At this stage, L4s were overlaid with 200 μl of *L. jeotgali* culture with an OD of 0.2-1.0. After 2 h of exposure to microbiome bacteria, the L4s were harvested with M9 + 0.05% Triton-X and washed twice in M9. Once washed, the animals were plated on new 10 cm NGM plates seeded with OP50-1. At 48 hours post exposure (hpe), the animals were harvested and washed again as before. Samples colonized with LUAb18 were directly imaged, while those with LUAb3 were stained using fluorescent in situ hybridization (FISH) probes using a LUAb3-specific probe, as described previously (Rivera et al. 2022; Poirier et al. 2024). The FISH probe (b003_16S_A) targets the 16S rDNA sequence CTCTCTGTGCTACCGCTCG of LUAb3, conjugated to either 5-Carboxyfluorescein (FAM) or CAL Fluor Red 610 (CF610). Imaging was performed using the fluorescent Eclipse Ni microscope (Nikon) at 20x magnification (Tran et al. 2022).

## Results

### Forward genetic screen for loss of bacterial binding in the gut

We previously described the discovery of a multiple *Lelliottia* spp. bacteria in the gut lumen of wild-caught *Caenorhabditis* spp., including *C. elegans* and *C. briggsae*. Focusing on *L. jeotgali* isolate LUAb3 and *C. elegans* N2 (wild-type strain), we demonstrated that this bacteria can actively replicate in the gut lumen and colonize the entire anterior-posterior axis of nearly 98% of a population of animals (Rivera et al. 2025). Additionally, LUAb3 colonization had no net negative effect on host fitness, allowing us to characterize LUAb3 as a commensal gut microbiome bacterium (Rivera et al. 2025). By Nomarski microscopy, we could visualize LUAb3 colonization as a continuous layer of bacilli that attach to the gut lining (Rivera et al. 2025). Given the capacity of LUAb3 to efficiently colonize N2 through attachment, we predicted that a forward genetic screen for loss of colonization would allow for the discovery of host genes required for bacterial binding in the gut.

To easily visualize colonization for a screen, we created a transgenic LUAb3 containing tdTomato inserted into the genome using mini-TN7 (Poirier et al. 2024). This *L. jeotgali*-tdTomato strain showed a similar attachment phenotype in the gut (Fig. 1A) and could colonize >98% of a population of an N2 population after just a 2 hr pulse exposure (Fig. 1B). Additionally, transformed LUAb18 (*L. jeotgali*-tdTomato) had no net negative effect on host lifespan (Fig. 1C), similar to LUAb3 (Rivera et al. 2025). Using this phenotype, we designed a forward genetic EMS screen to look for mutant animals that lost *L. jeotgali*-tdTomato colonization on a fluorescent dissecting microscope (Fig. 1D). We named this phenotype Adhering bacteria colonization deficient (Abcd). A total of three independent screens were conducted resulting the isolation of 42 self-propagating lines that had consistent, reproducible Abcd phenotypes (named Abcd1-Abcd42) (Fig. 1E). To ensure a true loss of LUAb3 colonization, fluorescent in situ hybridization (FISH) with a LUAb3-specific 16S rDNA probe was implemented on a subset of these lines to confirm the loss of LUAb18 colonization (Fig. 1F).

**Figure 1.**
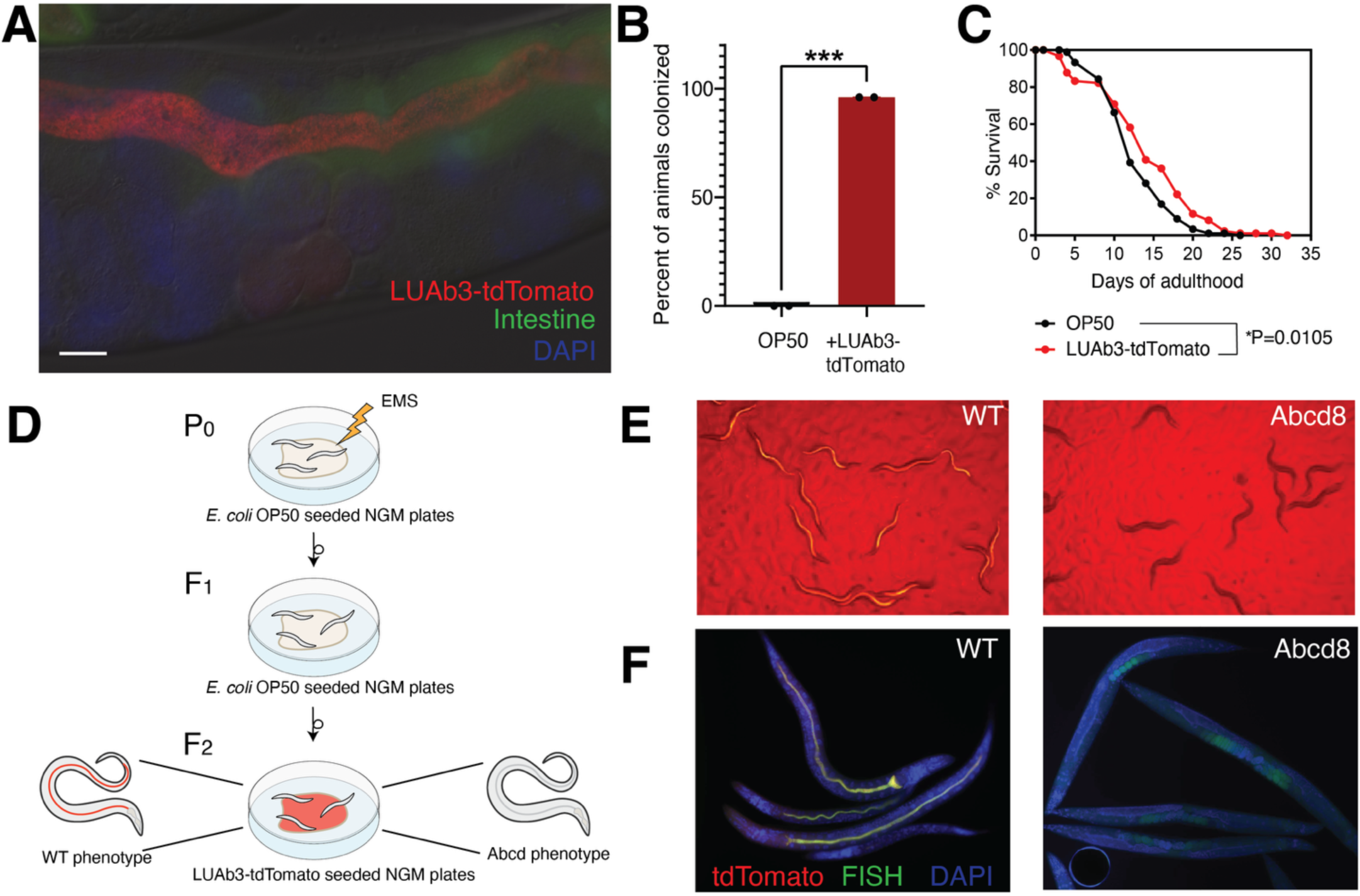
Host screen for loss of *L. jeotgali* adherence. (A) Fluorescence micrograph image of LUAb3-tdTomato colonizing *C. elegans* strain ERT413 expressing GFP in the intestine. Nuclei stained with DAPI. Scale bar is 10 microns. (B) Percentage of WT *C. elegans* N2 colonized with or without LUAb3-tdTomato in the intestine at 48 hpe (N = 2, n = 30). *** is p<0.005 by two-tailed t test. (C) Lifespan of N2 colonized with LUAb3-tdTomato. Sample size, n = 60 in N=3 independent experiments. Statistical analysis performed by Mantel-Cox test, where *p = 0.0105. (D) Schematic for the EMS mutagenesis screen used in this study to look for mutants that lose colonization by LUAb3-tdTomato. (E) Example image of mutant line Abcd8 (LUA66 *kola-*3 *ver21*) colonized with LUAb3-tdTomato in post screen validation. WT N2 is included as a positive control. L4s were colonized for 2 hours, then transferred to OP50 seeded NGM plates for 48 hours and imaged on the plate with an SMZ fluorescent microscope. (F) Fluorescence micrograph images of N2 and Abcd8 colonized as above in E, but then stained for LUAb3 with a FAM-conjugated FISH probe specific to the 16s rRNA of *L. jeotgali*.

### Identification of genes that regulate bacterial binding in the gut

To identify the genes associated with the Abcd phenotype, we conducted whole genome sequencing and in silico complementation of all mutant lines (Minevich et al. 2012). After subtracting all background single nucleotide variants found in our N2 strain and in >75% of all sequenced Abcd strains, we searched for genes with multiple moderate to strong alleles that affected the amino acid sequence of the protein in different mutants. Because each of the genes discovered in this manner were unclassified, we named them *kola* genes for *k*nock *o*ut of L*elliottia a*dherence. In this manner, we found 6 alleles in the gene K06A9.1 (*kola-1*), 16 in R08E3.1 (*kola-2*), 6 alleles in K06G5.1 (*kola-4*), and 5 alleles in C49C3.4 (*kola-5*). Separately, we found that two strains contained two likely null alleles (an early nonsense and splice site mutation) in C01B10.6 (*kola-3*). Four mutants that were not identified using in silico complementation and were backcrossed four times to N2 and re-subjected to WGS. Only 1 new gene was identified, *gly-5*, with a nonsynonymous G113R mutation *gly-5*(*ver34*). In this manner, all but two Abcd mutants were identified to have candidate alleles in these six genes (Table 2).

**Table 2.** Genes and alleles that result in a loss of *L. jeotgali* colonization.

| Gene symbol | Systematic name | Unique alleles | Strain names (*=backcrossed) | Mutation type | Chr | Chr Position | Mutation<sup>1</sup> | Amino acid change<sup>2</sup> | Phenotype |
|---|---|---|---|---|---|---|---|---|---|
| *kola-1* | K06A9.1 | *ver1* | LUA49 | missense | X | 1549060 | G7394A | G2465E | >90% abcd |
|  |  | *ver2* | LUA55 | missense | X | 1559625 | G7688A | G2563E | >90% abcd |
|  |  | *ver3* | LUA65* | splice donor | X | 1559075 | G>A | N/A | >90% abcd |
|  |  | *ver4* | LUA143* | missense | X | 1559515 | C7628T | T2543I | >90% abcd |
|  |  | *ver5* | LUA146* | missense | X | 1559192 | C7586T | A2529V | >90% abcd |
|  |  | *ver6* | LUA149* | splice acceptor | X | 1558454 | G>A | N/A | >90% abcd |
| *kola-2* | R08E3.1 | *ver7, ver8* | LUA52, LUA42 | splice acceptor | X | 4831828 | G>A | N/A | >90% abcd |
|  |  | *ver9* | LUA44 | splice acceptor | X | 4831649 | G>A | N/A | >90% abcd |
|  |  | *ver10* | LUA64* | missense | X | 4834101 | T5684A | L1895H | >90% abcd |
|  |  | *ver11* | LUA63* | missense | X | 4830554 | G4358A | C1453Y | >90% abcd |
|  |  | *ver12* | LUA46 | missense | X | 4830023 | G3916A | G1306R | >90% abcd |
|  |  | *ver13* | LUA39 | missense | X | 4833826 | G5504A | C1835Y | >90% abcd |
|  |  | *ver14, ver15* | LUA48, LUA71 | missense | X | 4829391 | G3476A | G1159E | >90% abcd |
|  |  | *ver16* | LUA142* | splice acceptor | X | 4827897 | G>A | N/A | >90% abcd |
|  |  | *ver17* | LUA79 | missense | X | 4833865 | T5543G | I1848S | >90% abcd |
|  |  |  |  | nonsense | X | 4833955 | T5586A | Y1862X |  |
|  |  | *ver19* | LUA147*` | nonsense | X | 4824284 | T764A | L255X | >90% abcd |
|  |  | *ver20* | LUA91 | missense | X | 4832166 | C4913T | T1638I | >90% abcd |
|  |  | *ver35* | LUA140* | deletion (124 bp) | X | 4831620-743 | N/A | N/A | >90% abcd |
|  |  | *ver36* | LUA148* | deletion (144 bp) | X | 4833074-217 | N/A | N/A | >90% abcd |
| *kola-3* | C01B10.6 | *ver21* | LUA66* | splice acceptor | IV | 6642501 | G>A | N/A | >90% abcd |
|  |  | *ver22* | LUA141* | nonsense | IV | 6641578 | C115T | Q39X | >90% abcd |
| *kola-4* | K06G5.1 | *ver23* | LUA132* | missense | X | 14217969 | T1038A | N346K | >90% abcd |
|  |  | *ver24* | LUA47 | missense | X | 14217297 | G722A | G241E | >90% abcd |
|  |  | *ver25* | LUA136* | missense | X | 14218936 | G1477A | G494R | >90% abcd |
|  |  | *ver26* | LUA114* | nonsense | X | 14218930 | C1471T | Q491X | >90% abcd |
|  |  | *ver27* | LUA115* | nonsense | X | 14217880 | C949T | Q317X | >90% abcd |
|  |  | *ver28* | LUA78 | missense | X | 14216467 | G416T | G139V | >90% abcd |
| *kola-5* | C49C3.4 | *ver29* | LUA50 | nonsense | IV | 17322810 | G1204T | E402X | >90% abcd |
|  |  | *ver30* | LUA137* | nonsense | IV | 17327296 | C3766T | Q1256X | >90% abcd |
|  |  | *ver31* | LUA80 | missense | IV | 17325117 | T2756G | M919R | >90% abcd |
|  |  | *ver32* | LUA144* | missense | IV | 17327955 | G4160A | R1387K | >90% abcd |
|  |  | *ver33* | LUA138* | splice acceptor | IV | 17326207 | G>A | N/A | >90% abcd |
| *gly-5* | Y39E4B.12 | *ver34* | LUA145* | missense | III | 13188527 | G337A | G113R | >90% partial abcd<sup>3</sup> |

<sup>1</sup>mutation positions represent the longest transcript for genes with multiple splice variants
<sup>2</sup>amino acid poisitions represent the longest transcript for genes with multiple splice variants
<sup>3</sup>partial abcd is discontinuous *L. jeotgali* colonization along the A-P axis of the gut lumen

### Description of alleles

For *kola-1* (K06A9.1), we obtained 6 independent alleles in the screen, including 4 missense and 2 splice site mutations. KOLA-1 is predicted to encode a membrane-bound mucin (Zugasti et al. 2016), with a transmembrane (TM) domain at the N-terminus, and multiple splice variants. The largest transcript, K06A9.1b, encodes for a 2596 AA protein. All mutations from the screen clustered towards the 3’ end of the gene (C-terminus), a region that is common between all seven predicted splice variants (Fig. 2A). The *kola-1(ver6)* splice acceptor allele has the most severe predicted effect, potentially deleting a large portion of the entire C-terminal region where the other allelic changes are clustered.

**Figure 2.**
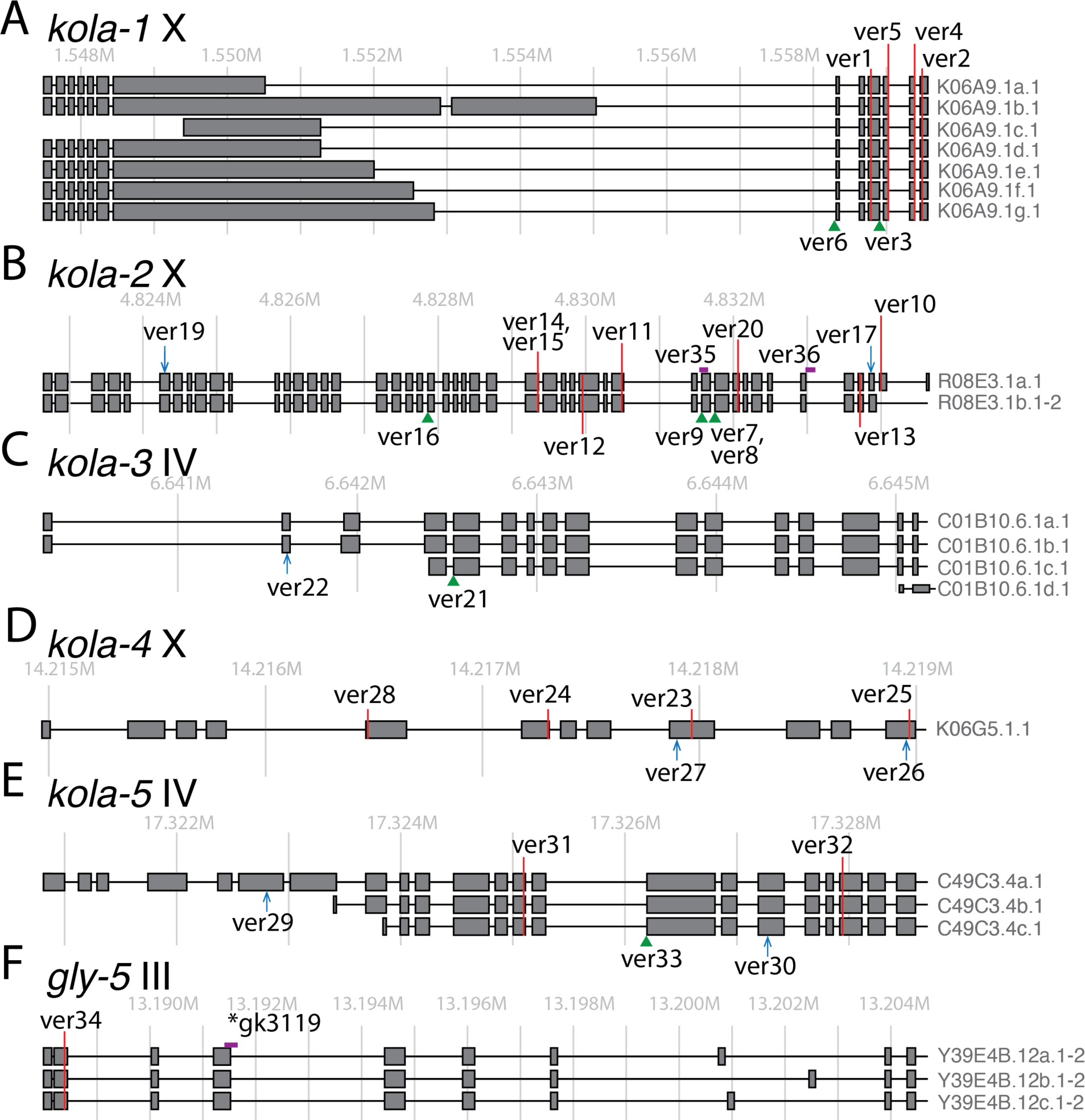
Gene model and map of alleles associated with loss of *L. jeotgali* binding. Each gene (A-F) and chromosome are indicated above the gene models, with estimated chromosomal locations marked by grey lines indicating 1 kilobase increments. Exons are indicated by dark gray boxes and introns by black lines connecting exons, with transcript isoform names on the right. Each *ver* allele from this screen are identified on the gene model with the approximate location of missense mutations indicated with a red line, nonsense mutations with blue arrows, splice site mutations with green arrowheads, and large deletions with magenta rectangles. Exons are grey boxes. \**gk3119* is an externally acquired *gly-5* mutant.

The *kola-2* (R08E3.1) gene was the most frequently hit gene in our screen with 13 unique alleles (Fig. 2B). Two lesions were each recovered in independent screens (*ver7*/*ver8* and *ver14*/*ver15*), which means that across the three screens we acquired 15 independent *kola-2* mutational events. The *kola-2* gene has no known prior role or function, but it is predicted to encode for a protein that enables Notch binding activity. The KOLA-2 protein has a predicted signal peptide (SP) but lacks a TM domain, indicating that the protein is likely secreted. The mutations span across the entire *kola-2* gene, with clustering occurring in the latter third, suggesting a mutational hotspot that affects *L. jeotgali* binding. In total, there were six unique missense mutations, three splice site mutations, two nonsense mutations, and two deletions over 100 bp in length. One allele, *ver19*, is a very early nonsense allele that likely results in a null, despite no obvious growth phenotype in the mutant strains. There are two predicted transcripts for this gene, but the *kola-2(ver10)* missense allele only effects the larger transcript R08E3.1a, suggesting that this splice variant alone is required for *L. jeotgali* colonization.

The *kola-3* (C01B10.6) gene was discovered in our screen because two different mutants contained high-impact alleles, *ver22* being an early nonsense allele, and *ver21* being an early splice acceptor allele (Fig. 2C). The *kola-3* gene encodes for a protein with no known prior role, but it has a predicted signal peptide (SP) and no TM domain, suggesting it is in the secretory pathway. Both alleles likely result in a null, despite no obvious growth phenotype in the mutant strains. The *kola-3(ver22)* allele only affects the two largest transcripts (C01B10.6.1a-b), suggesting the smaller transcript is not required for *L. jeotgali* colonization.

For *kola-4* (K06G5.1), we obtained 6 independent alleles, including 4 missense and two nonsense alleles (Fig. 2D). This gene had no known prior role or function but was predicted to encode a von Willebrand factor type A (VWFA) domain-containing protein. The KOLA-4 protein has a predicted signal peptide (SP) but lacks a TM domain. The mutations spanned across the latter two-thirds of the *kola-4* gene, with no obvious clustering.

For *kola-5* (C49C3.4), we obtained 5 independent alleles, including two missense, two nonsense, and one splice site mutation (Fig. 2E). Like *kola-2-4*, the *kola-5* gene had no known prior role or predicted function. Similarly, the KOLA-5 protein has a predicted signal peptide (SP) but lacks a TM domain. *kola-5* has three splice variants, but *ver29* is a nonsense allele found only in the largest transcript C49C3.4a.1, suggesting that only this transcript is required for *L. jeotgali* adherence.

The single *gly-5(ver34)* mutation found in this screen was a G113R missense mutation occurring early in the gene (Fig. 2F). *gly-5* encodes for a UDP-GalNAc:polypeptide N-acetylgalactosaminyltransferase (ppGaNTase) with a single-anchor type II membrane protein predicted to be localized to the Golgi apparatus (Hagen and Nehrke 1998; Hang and Bertozzi 2005). The location of the *gly-5(ver34)* mutation in the protein lies upstream of the two ppGaNTase catylase domains (AA174-407) and the ricin-like lectin domain (AA488-610) (Hagen and Nehrke 1998; Fritz et al.).

### Validation of alleles

To test whether independent *kola* alleles affect the same gene, we performed classical complementation tests, crossing hermaphrodites carrying one allele to males carrying another and scoring colonization in the F_1_. Crosses between two alleles of the same gene (*kola-1, kola-3* and *kola-4*) failed to complement, as F_1_ progeny remained Abcd (≤5% colonized; Fig. 3A). Crosses between alleles of different *kola* genes complemented, with 10–31% of F _1_ hermaphrodites colonized (Fig. 3A). This fraction is modest because *C. elegans* hermaphrodites self-fertilize, so many F1 hermaphrodites are self-progeny that retain the parent’s Abcd phenotype and only a subset are cross-progeny. F_1_ males are essentially all cross-progeny, so we also scored colonization in males. When F_1_ males were Abcd even though F_1_ hermaphrodites were colonized this indicates that the P_0_ hermaphrodite allele is X-linked (*kola-1(ver5)* and *kola-2(ver10)*), because males are hemizygous and inherit only X from the hermaphrodite.

**Figure 3.**
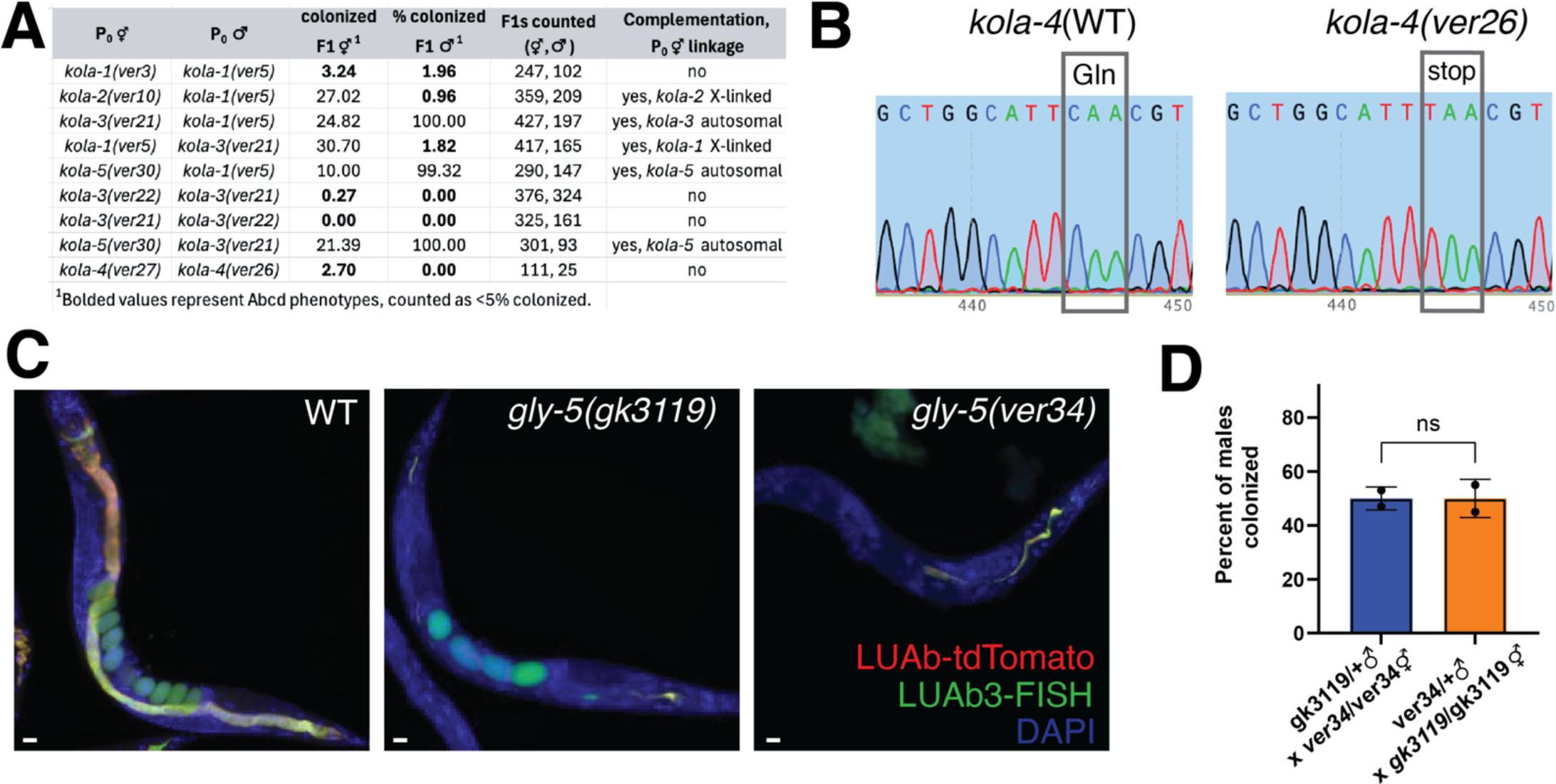
Validation of *kola* and *gly-5* mutants. (A) Complementation tests of *kola* alleles, with the indicated P_0_ alleles and scoring for F_1_ male and hermaphrodite populations for colonization by LUAb3-tdTomato at 48 hpe. (B) Example Sanger sequencing chromatographs of the lesion site of *kola-4*(*ver26*) nonsense mutation after PCR amplification, with WT N2 as a control. (C) Fluorescence micrograph images of N2, *gly-5*(gk3119), and *gly-5*(*ver34*). L4s were colonized for 2 hours, transferred to OP50 seeded NGM plates for 48 hours, and stained for LUAb3 with a FAM-conjugated FISH probe specific to the 16s rRNA of *L. jeotgali*. Scale bars are 20 microns. (D) Indirect complementation test for *gly-5* using outcrossed heterozygous males. Percentage of F_1_ males from the indicated cross colonized with LUAb3-tdTomato in the intestine at 48 hpe (N = 2, n = 30). ns is not significant by two-tailed t test.

For further validation, multiple alleles of each *kola* gene were backcrossed into N2 males four times (strains with an **\*** in Table 2, column 4). For each backcross, we selected F_1_ animals for gain of colonization with LUAb3-tdTomato to distinguish cross progeny. Then, F_2_s were selected for loss of colonization. After 4 consecutive backcrosses, strains were genotyped by PCR for the corresponding *kola* mutation and all tested mutants maintained their *kola* alleles (Fig. 3B). Note that all the EMS mutants used for complementation analysis were found to be recessive, as all F1 cross progeny from backcrossing exhibited a wild-type adherence phenotype.

We obtained only a single *gly-5* mutant in the screen, which displayed a partial Abcd phenotype, defined as discontinuous *L. jeotgali* colonization across the anterior-posterior axis of the lumen (Fig. 3C). To validate its role in adherence, we obtained an external allele, *gly-5*(*gk3119*), which has a deletion spanning from exon 4 to intron 4 of the gene (see Fig. 2F). We found that this strain also displayed a partial Abcd phenotype, similar to *gly-5*(*ver34*) (Fig. 3C). To further validate the role of *gly-5* in *L. jeotgali* adherence, we also conducted complementation. Unfortunately, both *gly-5* alleles failed to produce males using traditional heat shock. Therefore, we implemented an indirect test using outcrossed heterozygous males. For this, N2 males were crossed into both *gly-5* alleles. The resulting *gly-5* males were crossed into the hermaphordites homozygous for the other allele, i.e. *gly-5(ver34)/+* males x *gly-5(gk3119)/gly-5(gk3119)* hermaphrodites. The male progeny from both reciprocal crosses were tested for LUAb3-tdTomato colonization. These cross-progeny males from both crosses demonstrated an ∼50% rescue of the Abcd phenotype, indicative of a lack of complementation in *ver34/gk3119* trans-heterozygotes, whereas the other ∼50% of males were rescued by complementation from the WT *gly-5* allele (Fig. 3D). Note that only males were scored in this assay to eliminate scoring of any self-progeny from unmated *gly-5* hermaphrodites.

## Discussion

We describe the first unbiased, forward genetic screen to identify host factors required for commensal microbiome adherence. To conduct this screen, we utilized a genetically amenable microbiome bacterial species, *L. jeotgali*, with a distinct capacity to attach to the epithelial cells and actively replicate in the *C. elegans* gut lumen (Rivera et al. 2025). *L. jeotgali-*tdTomato made it easy to visualize colonization and we identified over 40 mutants with a near complete loss of bacteria in the gut lumen. In silico complementation was used to identify multiple alleles in five *kola* genes. Each of the *kola-1-5* genes were unannotated, supporting the notion that we hit new genes that play a vital role in *L. jeotgali* colonization.

A major goal of this screen was to identify a potential *L. jeotgali* receptor, and each of the *kola* genes may perform such a role, with *kola-1* being the top candidate as a putative membrane-bound mucin. What was unexpected from this screen was the identification of four different proteins predicted to be secreted (*kola-2-5*), each independently being required for bacterial colonization, suggesting a potentially complex mechanism for colonization. Another major goal of this screen was to identify signaling genes that regulate microbiome colonization and, surprisingly, we did not hit a single gene predicted to encode for a cytoplasmic protein or transcription factor. We have conducted three independent versions of this screen and given the number of times we hit *kola-2*, it is likely that the screens are near saturation. It is possible that there are other *C. elegans* genes that regulate bacterial adherence but were not viable. Another surprising aspect of the screen is that after backcrossing to N2, the most severe alleles of each *kola* gene were viable, with no obvious, superficial effects on growth, lifespan, or progeny. This suggests that disruption of the capacity for microbiome bacterial binding has no severe effect on development or fitness.

One additional gene we hit in our screen and validated with an independent mutant and complementation was *gly-5*. This gene encodes for a ppGaNTase, which initiates the biosynthesis of mucin-type O-linked glycosylation through addition of the N-acetylgalactosamine (GalNAc) to specific threonine and serine residues on proteins in the secretory pathway (Hang and Bertozzi 2005). Multiple splice forms of *gly-5* have displayed ppGaNTase enzymatic activity in vitro (Hagen and Nehrke 1998), and this gene was observed to be expressed in the intestine by both transcriptomics and proteomics (Kaletsky et al. 2018; Tan et al. 2024). The *gly-5(ver34)* G113R mutation occurs early in the gene and is predicted to affect all three transcript isoforms. While this allele is upstream of the known ppGaNTase catylase and ricin-like lectin domains in the protein, the glycine residue that is changed is a conserved residue among *C. elegans* ppGaNTase proteins (Hagen and Nehrke 1998). These data suggest the mucin-type O-linked glycosylation in the intestine may be required for *L. jeotgali* adherence in *C. elegans*.

Overall, we conducted a screen to understand the molecular basis of bacterial adherence in the *C. elegans* gut, especially in the context of an ecologically relevant *C. elegans* commensal. Given that forty percent of *C. elegans* genes lack a functional annotation, it is possible that identification of the function of many genes requires a natural context, including growth with a natural microbiome.

## Data availability

All data necessary for confirming the conclusions of the article are present within the article, figures, tables, and supplements. *C. elegans* and bacterial strains are available upon request.

